# The Development of Cognitive Control in Children with Autism Spectrum Disorders or Obsessive-Compulsive Disorder: A Longitudinal fMRI study

**DOI:** 10.1101/2020.04.09.033696

**Authors:** Bram Gooskens, Dienke J. Bos, Jilly Naaijen, Sophie E.A. Akkermans, Anna Kaiser, Sarah Hohmann, Muriel M.K. Bruchhage, Tobias Banaschewski, Daniel Brandeis, Steven C.R. Williams, David J. Lythgoe, Jan K. Buitelaar, Bob Oranje, Sarah Durston, the TACTICS consortium

**Affiliations:** Department of Psychiatry, Brain Center, University Medical Center Utrecht, Utrecht University, Utrecht, The Netherlands; Donders Centre for Cognitive Neuroimaging, Donders Institute for Brain, Cognition and Behaviour, Radboud University, the Netherlands; Department of Cognitive Neuroscience, Donders Institute of Brain, Cognition and Behaviour, Radboud University Medical Center, Nijmegen, The Netherlands; Department of Child and Adolescent Psychiatry and Psychotherapy, Central Institute of Mental Health, Medical Faculty, Mannheim/Heidelberg University, Mannheim, Germany; Department of Neuroimaging, King’s College London, Institute of Psychiatry, Psychology and Neuroscience, London, UK; Department of Child and Adolescent Psychiatry and Psychotherapy, Psychiatric Hospital, University of Zurich, Zurich, Switzerland; Center for Integrative Human Physiology, University of Zurich, Zurich, Switzerland; Neuroscience Center Zurich, University of Zurich, Zurich, Switzerland; ETH Zurich, Zurich, Switzerland; Karakter Child and Adolescent Psychiatry University Center, Nijmegen, The Netherlands

**Keywords:** Autism Spectrum Disorders, Obsessive-Compulsive disorder, repetitive behavior, cognitive control, fMRI

## Abstract

Repetitive behavior is a core symptom of Autism Spectrum Disorder (ASD) and Obsessive-Compulsive Disorder (OCD), and has been associated with impairments in cognitive control. However, it is unclear how cognitive control and associated neural circuitry relate to the development of repetitive behavior in children with these disorders. In a multicenter, longitudinal study (TACTICS; Translational Adolescent and Childhood Therapeutic Interventions in Compulsive Syndromes), the development of cognitive control was assessed during late childhood using a longitudinal fMRI design with a modified stop-signal task in children with ASD or OCD, and typically developing (TD) children (baseline: N=122 (8-12y), follow-up: N=72 (10-14y), average interval: 1.2y). Stop-signal reaction time (SSRT) decreased over development, regardless of diagnosis. Repetitive behavior in children with ASD and OCD was not associated with performance on the stop-signal task. There were no whole-brain between-group differences in brain activity, but ROI-analyses showed increases in activity in right precentral gyrus over development for children with OCD. In sum, even though subtle differences were observed in the development of brain activity in children with OCD, the findings overall suggest that the development of cognitive control, as assessed by the stop signal task, is similar in children with and without ASD or OCD.

## 1. Introduction

Cognitive control is the ability to stop or suppress ongoing behavior when it is no longer appropriate or required. This ability is crucial for successfully navigating the demands of daily life. Although cognitive control starts to early early in development, it continues to develop into adulthood (Luna, Padmanabhan & O’Hearn, 2010; Luna et al. 2015). Impairments in cognitive control are thought to play a role in repetitive behavior and the emergence of neurodevelopmental disorders where such behaviors play a role, such as autism spectrum disorders (ASD) and obsessive-compulsive disorder (OCD) (Chamberlain et al., 2005; Hill, 2004; Lipszyc and Schachar, 2010; Moritz et al., 2002; Mosconi et al., 2009; Snyder et al., 2015).

Repetitive behaviors in ASD and OCD show qualitative and quantitative similarities. For instance, restricted interests in ASD and obsessions in OCD both involve persistent repetitive thoughts, whereas stereotyped behavior in ASD and compulsions in OCD both reflect symptoms that manifest as repetitive behavior that must be carried out (Jiujas et al., 2017). Furthermore, children with ASD and OCD have been reported to display similar levels of sameness- and repetitive sensory-motor behaviors (Zandt et al., 2007). Previous studies have demonstrated increased rates of obsessive-compulsive symptoms in ASD and vice versa (Ivarsson and Melin, 2008; Leyfer et al, 2006). However, the purpose of these behaviors or the context in which they take place may differ between the disorders. While repetitive behavior in ASD is typically maintained by reinforcement, in OCD it usually has the purpose of relieving anxiety or distress caused by the obsession (McDougle et al., 1995; Zandt et al., 2007; 2009). The extent to which ASD and OCD have distinct versus common neural correlates of repetitive behavior, and how this relates to the development of cognitive control is still unknown.

While cognitive control typically improves with development (Huizinga et al., 2006; Luna, Padmanabhan & O’Hearn, 2010; Luna et al., 2015), reports on age-related changes in cognitive control in ASD have been mixed. Some studies have shown no improvement or slight worsening (Rosenthal et al., 2013; Solomon et al., 2008), whereas others have suggested developmental improvements (Christ et al., 2011; Happe et al., 2006; Luna et al., 2007). In OCD, there have been only cross-sectional studies of cognitive control to date, and those have suggested that impairments in cognitive control are not as evident in children and adolescents (Gooskens et al., 2019; Marzuki et al., 2020; Rubia et al., 2010; Wooley et al., 2008) as in adults with OCD, where longer stop-signal reaction times (SSRT) have often been reported (Chamberlain et al., 2007; Kang et al., 2012; Penadés et al., 2007; de Wit et al., 2012).

Most functional Magnetic Resonance Imaging (fMRI) studies on the typical development of cognitive control have found increases in activity of prefrontal cortex (PFC) with development, often related to improvements in task performance (Bunge et al., 2002; Cohen et al., 2010; Durston et al., 2002; Rubia et al., 2007). However, age-related decreases in PFC activity have also been reported (Somerville et al., 2011), and have been suggested to reflect a developmental decrease in the effort required to exert cognitive control (Tamm et al., 2002), consistent with a shift from diffuse to focal engagement of PFC (Durston et al., 2006). Additionally, nonlinear developmental changes, where brain activity in PFC increases from childhood to adolescence, followed by decreases into adulthood have been reported (Luna et al., 2001; Somerville et al., 2011).

fMRI studies using the stop-signal task to compare brain activity in children with OCD to typically developing children have reported hypoactivation in brain regions associated with cognitive control, including in PFC regions such as the dorsolateral prefrontal cortex (DLPFC) and inferior frontal gyrus (IFG), but also striatum and thalamus (Carlisi et al., 2017; Rubia et al., 2010; Wooley et al., 2008). Contrary to hypoactivation observed in children with OCD, hyperactivation in PFC regions including the middle frontal gyrus (MCG) and the IFG has been reported in children with ASD during cognitive control (Carlisi et al., 2017; Chantiluke et al., 2015; Albajara Sáenz et al., 2020). However, longitudinal work investigating development of cognitive control and its neurobiological correlates is lacking.

In the current study, we recruited children with ASD, OCD and typically developing children (9-14 years) in the context of a multi-center collaborative initiative, the Translational Adolescent and Childhood Therapeutic Interventions in Compulsive Syndromes (TACTICS) consortium. The primary objective of this study was to investigate the development of cognitive control and associated neural circuitry in relation to repetitive behavior in children with ASD and OCD. We operationalized cognitive control as the ability to stop an ongoing response in the context of a stop-signal task during fMRI. At baseline (mean age = 10 years), we found no differences in cognitive control and associated neural circuitry in children with ASD or OCD compared to typically developing (TD) children. However, we did find that increased activity in prefrontal cortex was associated with ADHD symptoms (Gooskens et al., 2019). As cognitive control improves over development, and given that deficits in cognitive control in OCD may emerge later in development, we performed a follow-up study. We hypothesized that SSRT would decrease over development for all children, indicating improvement in cognitive control. We further hypothesized that developmental improvements in task performance would be associated with increased brain activity in prefrontal brain areas for typically developing children. Finally, we hypothesized that children with OCD and ASD would show delay in the development of cognitive control, reflected by smaller improvements in task performance and an attenuated increase in brain activity in prefrontal brain areas over development compared to TD children, and that these changes in cognitive control would relate to the presence and severity of repetitive behavior.

## 2. Methods

### 2.1 Participants

At baseline, we were able to include data for 122 participants between 8 and 12 years (ASD = 38, OCD = 23, TD = 61) (details are described in Gooskens et al., 2019). Ninety-three participants (ASD = 34, OCD = 15, TD = 44) participated in a follow-up visit. Participants’ reasons for drop-out were mainly loss of interest, or wearing dental braces which prevented them for undergoing MRI assessment. Average follow-up time was 1.23 years. Participants were seen at the same four sites across Europe (Central Institute of Mental Health, Medical Faculty Mannheim, University of Heidelberg Mannheim, Germany; King’s College London, London, United Kingdom; Radboud University Medical Center and the Donders Institute for Brain, Cognition and Behaviour, Nijmegen, The Netherlands; University Medical Center Utrecht Brain Center, Utrecht, The Netherlands) and were commissioned by a multicenter study (COMPULS: Naaijen et al., 2016) as part of the overarching TACTICS collaborative initiative (http://www.tactics-project.eu). Exclusion criteria for all participants were an estimated total IQ below 70 and insufficient comprehension and speaking ability of the native language of the country. For MR scanning, the presence of metal objects in the body (i.e., pacemaker, dental braces), neurological illness or other contra-indications for MRI-assessment were exclusion criteria. Participants were asked to abstain from stimulant medication 24 hours before scanning. For both diagnostic groups, a concurrent diagnosis of the other disorder was an exclusion criterion (i.e. comorbid OCD for a child with ASD, or vice versa). In TD participants or their first-degree relatives the presence of any psychiatric diagnosis was an exclusion criteria. After description of the study, parents of all participants gave written informed consent.

### 2.2 Phenotypic Information

Participants with ASD or OCD were diagnosed according to *The Diagnostic and Statistical Manual of Mental Disorders*, 4^th^ edition, Text Revision (APA, 2000) or 5^th^ edition criteria (APA, 2013). At baseline, ASD diagnosis was ascertained by a trained psychologist at each site using the Autism Diagnostic Interview-Revised (ADI-R; Lord et al., 1994). The Children’s Yale-Brown Obsessive-Compulsive Scale (CY-BOCS; Scahill et al., 1997) was used as a severity scale for obsessions and compulsions for all children with OCD. This interview was repeated at follow-up and also performed in participants with ASD if screening questions suggested the presence of clinically significant obsessions or compulsions.

To determine the presence of possible comorbidities, all parents were interviewed using the structured Diagnostic Interview Schedule for Children (DISC-IV, parent version; Shaffer et al., 2000), the Development and Well-being Assessment (DAWBA; Goodman et al., 2000) or the Kiddie Schedule for Affective Disorders and Schizophrenia (K-SADS; Kaufman et al., 1997) at both baseline and follow-up. Total Intellectual Quotient (IQ) was estimated using a shortened version of the Wechsler Intelligence Scale for Children (WISC-III; Wechsler, 2003). At both timepoints, repetitive behavior was assessed using the Repetitive Behavior Scale – Revised (RBS-R; Bodfish et al., 2000). In addition, the Conners’ Parent Rating Scale – Revised (CPRS-R:L; Conners, 2000) was used to rate Attention Deficit/Hyperactivity Disorder (ADHD) symptoms at both timepoints.

Information on medication was collected through parental report. Four children with ASD were being treated with psychostimulants, one with antipsychotics, one with a combination of both and one child with an antidepressant. Two children with OCD were being treated with psychostimulants, three with antidepressants, and one with antipsychotics and antidepressants. Three children with ASD had a current comorbid diagnosis of ADHD, one child had comorbid depressive disorder, and one child had both comorbid ADHD and oppositional defiant disorder (ODD). In the OCD group, one child had comorbid ADHD.

### 2.3 Stop-signal task

Our modified version of the stop-signal task (Rubia et al., 2003; Gooskens et al., 2019) requires participants withholding a motor response to a go stimulus when it is randomly followed by a stop signal. During go-trials (80% of trials), subjects were instructed to make a button response with their right index- or middle finger corresponding to the arrow direction (left arrow: index finger, right arrow: middle finger, duration arrow: 500 ms). The mean inter-trial interval (ITI) was randomly jittered between 1.6 and 2.0 s to optimize statistical efficiency. During stop trials (20% of trials), go-signals were followed (approximately 250 ms later) by arrows pointing upwards (stop signals), and participants were instructed to withhold (stop) their motor response. The delay between a go- and stop-signal (stop-signal delay: SSD) was dynamically adapted (start: 250 ms, step size: 50 ms) in response to each subject’s stop performance using a staircase algorithm. This procedure ensured that the session concluded with an approximately equal number of successful and failed stop-trials. Before scanning, participants performed a brief practice session of the task.

### 2.4 Task performance measures

The measures of interest for task performance were mean reaction time (MRT) on correct go-trials, stop-signal reaction time (SSRT), mean stop-signal delay (SSD), number of non-responses to go-trials (omissions) and number of incorrect responses to go-trials (commissions). The SSRT was estimated using the integration method from Verbruggen and colleagues (2013; 2019): first, reaction times (RT) to correct go-trials were rank ordered. Subsequently, the *n*th go-RT was selected, where *n* was derived by multiplying the number of correct go-trials by the probability that one would respond to a stop-signal (*P*(respond | stop-signal)). The SSRT was then estimated by subtracting the mean SSD from the *n*th go-RT.

### 2.5 Behavioral analysis

Baseline data that was excluded from statistical analysis is described in our previous paper (Gooskens et al., 2019). Of the ninety-six participants who participated in the follow-up visit, data from six participants could not be analyzed due to incomplete task performance (ASD N = 3, TD N = 3). To optimize data quality for statistical analysis, we used Congdon and colleagues’ method (2012) to exclude participants with SSRT values below 50 ms (ASD N = 6, OCD N = 1, TD N = 7) and accuracy below 25% on successful stop-trials (OCD N = 1) (Gooskens et al., 2019). As such, data were available for behavioral analysis for 122 participants at baseline (ASD N = 38, OCD N = 23, TD N = 61) and 72 participants at follow-up (ASD N = 25, OCD N = 13, TD N = 34) (see Table 1 and 2 for sample characteristics).

**Table 1.**
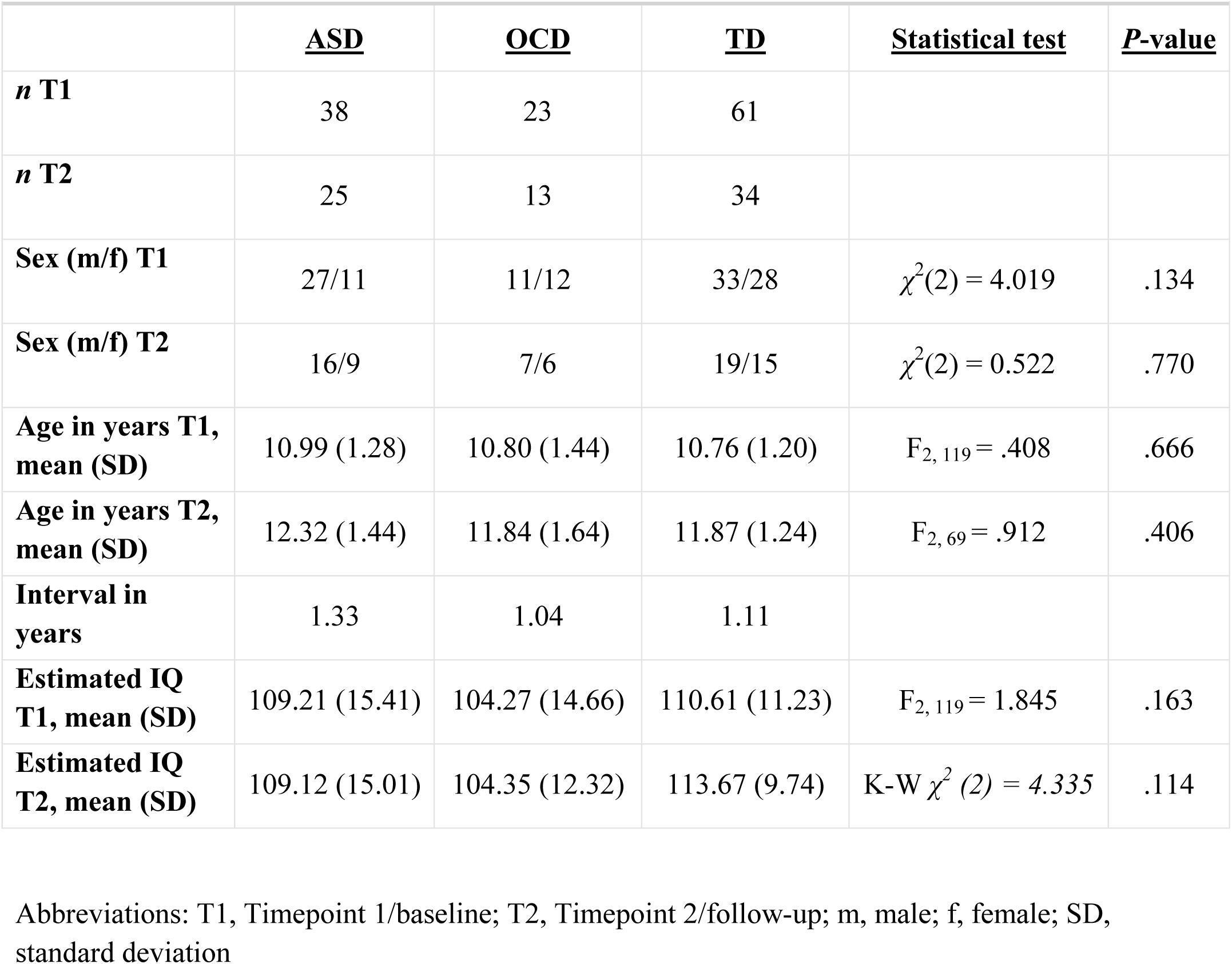
Sample characteristics [full sample].

|  | <u>**ASD**</u> | <u>**OCD**</u> | <u>**TD**</u> | <u>**Statistical test**</u> | <u>**P-value**</u> |
| --- | --- | --- | --- | --- | --- |
| ***n* T1** | 38 | 23 | 61 |  |  |
| ***n* T2** | 25 | 13 | 34 |  |  |
| **Sex (m/f) T1** | 27/11 | 11/12 | 33/28 | $\chi^2(2) = 4.019$ | .134 |
| **Sex (m/f) T2** | 16/9 | 7/6 | 19/15 | $\chi^2(2) = 0.522$ | .770 |
| **Age in years T1, mean (SD)** | 10.99 (1.28) | 10.80 (1.44) | 10.76 (1.20) | $F_{2, 119} = .408$ | .666 |
| **Age in years T2, mean (SD)** | 12.32 (1.44) | 11.84 (1.64) | 11.87 (1.24) | $F_{2, 69} = .912$ | .406 |
| **Interval in years** | 1.33 | 1.04 | 1.11 |  |  |
| **Estimated IQ T1, mean (SD)** | 109.21 (15.41) | 104.27 (14.66) | 110.61 (11.23) | $F_{2, 119} = 1.845$ | .163 |
| **Estimated IQ T2, mean (SD)** | 109.12 (15.01) | 104.35 (12.32) | 113.67 (9.74) | $K-W \chi^2(2) = 4.335$ | .114 |

Abbreviations: T1, Timepoint 1/baseline; T2, Timepoint 2/follow-up; m, male; f, female; SD, standard deviation

**Table 2.** Clinical measures and task performance [full sample].

|  | T1 | T2 | T1 | T2 | T1 | T2 | Statistics |  |  |
| --- | --- | --- | --- | --- | --- | --- | --- | --- | --- |
|  | <u>ASD</u><br>N = 38 | <u>ASD</u><br>N = 25 | <u>OCD</u><br>N = 23 | <u>OCD</u><br>N = 13 | <u>TD</u><br>N = 61 | <u>TD</u><br>N = 34 | <u>Time</u> | <u>Group</u> | <u>Time*</u><br><u>Group</u> |
|  | Mean (SD) | Mean (SD) | Mean (SD) | Mean (SD) | Mean (SD) | Mean (SD) | P-value | P-value | P-value |
| **<u>Questionnaires</u>** |  |  |  |  |  |  |  |  |  |
| *CY-BOCS* |  |  |  |  |  |  |  |  |  |
| - Obsessions |  |  | 7.00 (5.41) | 8.50 (5.41) |  |  | .839 | n.a. | n.a. |
| - Compulsions |  |  | 9.91 (4.02) | 8.62 (4.02) |  |  | .217 | n.a. | n.a. |
| - Total score |  |  | 16.48 (7.56) | 15.15 (10.23) |  |  | .317 | n.a. | n.a. |
| *RBS-Revised* |  |  |  |  |  |  |  |  |  |
| - Compulsivity | 2.68 (3.39) | 2.88 (4.46) | 4.65 (3.54) | 4.15 (4.31) | 0.05 (0.28) | 0.09 (0.29) | .210 | > .001*<sup>c</sup> | n.s. |
| - Total score | 23.32 (18.90) | 19.44 (18.45) | 17.52 (13.17) | 14.62 (11.23) | 0.56 (1.23) | 0.79 (1.97) | .864 | > .001*<sup>d</sup> | n.s. |
| *CPRS-Revised: Long<sup>f</sup>* |  |  |  |  |  |  |  |  |  |
| - Inattention | 62.52 (11.72) | 60.24 (10.34) | 57.36 (10.61) | 56.50 (14.22) | 45.83 (6.70) | 46.18 (6.05) | .817 | > .001*<sup>d</sup> | n.s. |
| - Hyperactivity | 62.90 (13.16) | 61.46 (13.62) | 61.29 (10.51) | 61.17 (16.18) | 46.12 (3.99) | 46.44 (4.93) | .600 | > .001*<sup>d</sup> | n.s. |
| - Total score | 64.40 (12.78) | 61.88 (11.77) | 59.93 (10.77) | 59.08 (14.35) | 45.44 (5.11) | 45.25 (4.61) | .661 | > .001*<sup>d</sup> | n.s. |
| **<u>Performance</u>** |  |  |  |  |  |  |  |  |  |
| MRT | 569.82 (79.84) | 508.35 (62.91) | 551.61 (77.33) | 558.34 (104.66) | 564.65 (81.91) | 546.92 (66.15) | .213 | .887 | n.s. |
| SSD | 350.94 (97.81) | 334.22 (72.70) | 317.98 (114.98) | 402.17 (98.18) | 345.10 (87.95) | 352.70 (65.68) | .067 | .781 | .022*<sup>b</sup> |
| SSRT | 186.30 (82.88) | 145.62 (48.74) | 211.34 (114.45) | 122.75 (54.54) | 186.70 (69.20) | 164.66 (59.83) | .009*<sup>a</sup> | .964 | n.s. |
| Omissions<sup>g</sup> | 5.22 | 2.35 | 2.11 | 1.58 | 2.26 | 1.96 | .535 | .007*<sup>e</sup> | n.s. |
| Comissions<sup>g</sup> | 4.84 | 3.94 | 4.33 | 3.09 | 4.29 | 4.20 | .685 | .682 | n.s. |
| Successful Stopping | 51.45 | 51.33 | 50.65 | 52.56 | 51.53 | 51.47 | .501 | .956 | n.s. |

Abbreviations: T1, Timepoint 1, T2, Timepoint 2, ASD, autism spectrum disorder; OCD, obsessive-compulsive disorder; TD, typically developing group; SD, standard deviation; m/f, male/female; ADI, Autism Diagnostic Interview; CY-BOCS, Children's Yale-Brown Obsessive-Compulsive scale; RBS, Repetitive-Behavior scale; CPRS, Conners' Parent Rating scale, MRT = Mean reaction time, SSD = Stop-signal delay, SSRT = Stop-signal reaction time, n.a. not applicable, n.s. not significant (removed from model)
<sup>a</sup> SSRT decreased over development across all groups
<sup>b</sup> SSD increased over development for children with OCD, compared to both children with ASD and TDC
<sup>c</sup> OCD > ASD > TD
<sup>d</sup> ASD, OCD > TD
<sup>e</sup> ASD > TD
<sup>f</sup> Displayed scores are based on T-scores
<sup>g</sup> Commissions, omissions and successful stopping are displayed in % of total trials

Behavioral data analysis was performed in SPSS version 25 (IBM) and R-software 3.5.1 (R Core Team, 2019). Baseline group differences in demographic and clinical measures were tested using the appropriate Pearson’s χ^2^-tests or one-way analyses of variance (ANOVA). We used Levene’s test for homogeneity of variance and Shapiro-Wilk normality tests to check if assumptions of homogeneity of variance and normality were met. If data did not meet these assumptions, we ran non-parametric tests (Kruskal Wallis rank sum test or Mann–Whitney U-test).

To examine whether there were any group differences in the developmental trajectory of task-performance, we performed an intent-to-treat analysis using linear mixed-effects (LME) models in the lme4 package in R (Bates et al., 2015). For each task performance measure of interest (MRT, SSD, SSRT, number of omission and commission errors) we fit LME models with diagnostic group, time point (1 and 2), and age as fixed factors, and within subject dependence as a repeated measures random factor. Age was included as a fixed factor in all analyses, given the importance of developmental stage when investigating growth trajectories. Given that there were no significant differences in sex between the diagnostic groups, this factor was subsequently left out of the model. Because site is not a systematic and reliable predictor for explaining relationships with other dependent variables, it was only included in the model as a covariate when its effect reached statistical significance. If the diagnostic status by time interaction did not render any significant effects, it was removed from the design. For significant interaction effects, we ran post-hoc pairwise comparisons using least-squares means.

In order to examine tentative associations between change in task performance and change in symptom severity, we calculated delta scores for participants with complete data (time 2 – time 1) for the task performance measures and questionnaire scores (RBS-R compulsive subscale, total score; CY-BOCS total score; CPRS-R total score). Subsequently, we ran Spearman’s correlations to relate change in performance to change in behavior. The number of administered CY-BOCS in the ASD group was too low (N = 6 at follow-up) to include in any further analyses.

Finally, as children with ASD and OCD often have symptoms of ADHD, and to replicate findings from our baseline paper (Gooskens et al., 2019), we repeated the analysis on ADHD symptomatology, where we used a median split to create two groups based on CPRS-R score (Low < 56; High > 57).

### 2.6 MRI scanner information

At the four different sites, comparable 3-Tesla MRI scanners were used (Siemens Trio and Siemens Prisma, Siemens, Erlangen, Germany; General Electric MR750, GE Medical Systems, Milwaukee, WI, USA; Philips 3T Achieva, Philips Medical Systems, Best, The Netherlands). We used classic, gradient echo EPI sensitive to BOLD MR contrast (TR: 2070ms, TE: 35ms). More detailed information on the scanners and sequences is available in the baseline paper (Gooskens et al., 2019).

Before participating in the MR session, children at each site were prepared for scanning using a simulation session with a mock scanner (except for Mannheim). In this session, children were familiarized with MR sounds, the button box needed for task completion, and lying still in the scanner environment. If a child (or his/her parent) reported enhanced anxiety to enter the MR scanner, the session was ended. This procedure has proven successful in reducing anxiety for the MR session (Durston et al., 2009).

### 2.7 fMRI preprocessing

fMRI preprocessing was performed using standard procedures in SPM12, as implemented in MATLAB R2015b. fMRI data were realigned to the first volume to correct for in-scanner head motion. Next, using the ArtRepair toolbox in SPM12, all volumes with frame-to-frame movement >1 mm or >1.5% standard deviation from the mean signal were substituted using linear interpolation from neighboring frames. After realignment and motion-correction, the fMRI data and anatomical T1-image were co-registered, followed by normalization to Montreal Neurological Institute (MNI) standard atlas and finally spatially smoothed with a 6mm^3^ full width at half maximum (FWHM) Gaussian kernel.

### 2.8 fMRI data analyses

Of the data from 72 participants who successfully performed the fMRI stop-signal task at follow-up, fourteen datasets were excluded due to excessive head motion (> 3mm absolute movement) (ASD N = 3, OCD N = 4, TD N = 2) or replacement of more than 20% of total volumes (ASD N = 1, OCD N = 1, TD N = 3), both measured in the ArtRepair step. This resulted in 26 ASD, 16 OCD and 53 TD baseline datasets, and 21 ASD, 9 OCD and 29 TD follow-up datasets to carry forward to the fMRI analysis (see Supplement Table S2 and S3 for sample characteristics). For details about fMRI data exclusions at baseline we refer to our earlier paper (Gooskens et al., 2019).

At the first level, onsets of three event types (correct go-trials, successful stop-trials, failed stop-trials) were modeled using delta functions convolved with the canonical haemodynamic response function (HRF). Six motion estimation parameters were included in the model as regressors of no interest.

Second-level random effects analyses were run for two contrasts of interest: successful stopping was investigated by contrasting successful stop trials with correct go trials (successful stop activation > go activation). Failed stopping was investigated by contrasting failed stop trials with correct go trials (failed stop activation > go activation).

In the full fMRI sample, developmental differences between diagnostic groups were assessed with LME models, using data-driven ROIs that were defined in the baseline study (Gooskens et al., 2019). For each ROI, LME models were fit with diagnostic status, time point, and age as fixed factors, and within subject dependence as a repeated measures random factor.

To confirm that no bias was introduced by the participants with data at only one of the timepoints, the LME-analyses were repeated in a subsample of participants who had complete data at both timepoints (N = 40, sample characteristics are in Supplemental Table S4 and S5). Further, whole-brain main- and interaction effects were explored for the two contrasts (successful/failed stopping) using the ‘flexible factorial’ module in SPM12. To increase power, children with ASD (N = 14) and OCD (N = 6) were combined into a single group, similar to our previous paper. Subsequently, a design matrix was created that included the factor ‘subject’, the group variable ‘diagnostic status’ (ASD-OCD/control) and the within-subject factor ‘time’ (timepoint 1/timepoint 2). All tests were Family-Wise Error (FWE) corrected with a p-value of .05.

We assessed possible correlations between change in brain activity (whole brain and ROI) and change in task performance and symptom severity. We used the delta scores (time 2 – time 1) for the ROIs, task performance and questionnaires (RBS-R compulsive subscale, total score; CY-BOCS total score; CPRS-R total score) and calculated Spearman’s correlations. *P*-values were adjusted for multiple testing, using the Benjamini-Hochberg procedure to control the False Discovery Rate (FDR) (Benjamini & Hochberg, 1995).

## 3. Results

### 3.1 Group characteristics

The characteristics of the full (behavioral) sample are provided in Table 1. For this sample, diagnostic groups did not differ in age, sex, or estimated IQ. Children with ASD or OCD scored higher on the compulsivity subscale and total score of the RBS-R, and on the CPRS-R compared to typically developing children (Table 2). Moreover, children with OCD scored higher on the compulsivity scale of the RBS-R than children with ASD. In each diagnostic group, drop-outs did not differ in age, IQ, symptomatology or in any of the task performance measures, compared to the children that were included in the study, indicating bias due to drop-out is unlikely.

The fMRI subsample did not differ in age or sex, similar to the full sample. However, IQ differed between groups at baseline, with lower scores for children with OCD than typically developing children (see Supplemental Table S1). For children with OCD, compulsive behavior on the CY-BOCS and the RBS-R compulsivity scale decreased with development, whereas for children with ASD the RBS-R total score decreased with development. Similar to the full sample, children with ASD or OCD had significantly higher scores on the compulsivity scale and total score of the RBS-R, and on the CPRS-R than TD children, in the fMRI sample (Supplemental Table S2).

To confirm that no bias was introduced by participants with data at only one timepoint, we repeated the analyses in the subsample of participants who had complete data at both timepoints (N = 40). Characteristics of this study-completer sample are given in Supplemental Table S3.

### 3.2 Behavioral effects

Linear mixed-effects (LME) models showed a main effect of development (time 1 > time 2) [*F*_1,155_ = 6.825, *p* = .009] and an effect of age [*F*_1,122_ = 4.037, *p* = .046] on stop-signal reaction time (SSRT) in the full sample (Fig. 1, Table 2). In the fMRI sample, we found a main effect of development on SSRT only (time 1 > time 2) [*F*_1,117_ = 4.028, *p* = .047] (Fig.1, Table S2). We found no group differences in SSRT at baseline or follow-up. Yet, we found an interaction effect between diagnostic group and development on SSD in both samples (full: [*F*_2,104_ = 3.809, *p* = .022]; fMRI: [*F*_2,105_ = 3.114, *p* = .049]), where SSD increased over development for children with OCD, but not for children with ASD and TD children. In addition, we found an age effect on mean reaction time (MRT) in the full [*F*_1,137_ = 9.165, *p* = .003] and fMRI sample [*F*_1,146_ = 7.099, *p* = .009], indicating that reaction time decreased with age. Finally, we found an interaction effect between diagnostic group and development on MRT in the fMRI sample only [*F*_2,146_ = 5.287, *p* = .006], where reaction time decreased over development for children with OCD and TD children, but not children with ASD.

**Fig 1.**
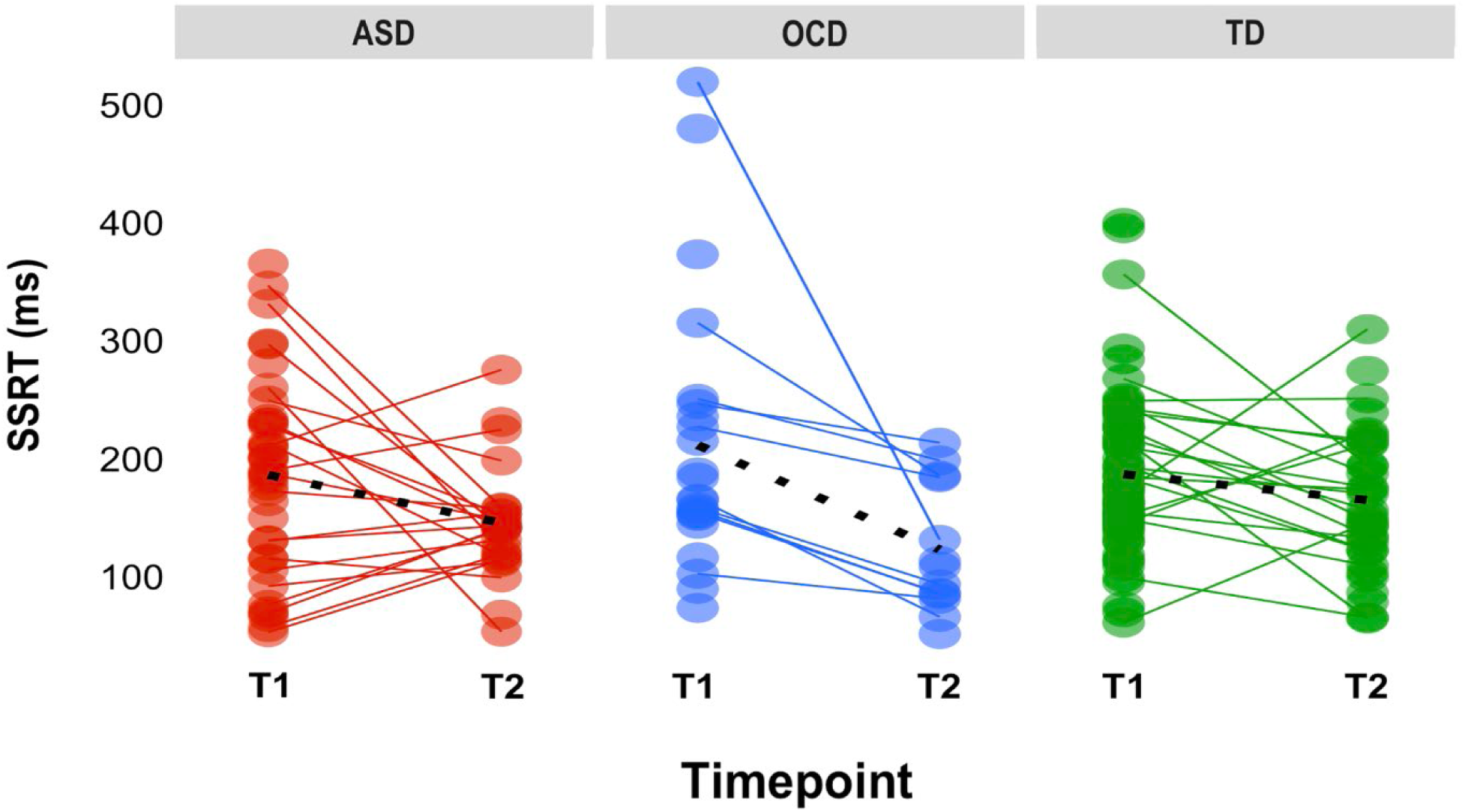
Stop-signal reaction time (SSRT) scores at baseline and follow-up for the three diagnostic groups. Black dashed line represents average slopes. SSRT decreased over development for all children, regardless of diagnosis. Abbreviations: ASD, Autism Spectrum Disorder; OCD, Obsessive-Compulsive Disorder; TD, typically developing; ms, milliseconds; T1, timepoint 1/baseline; T2, timepoint 2/follow-up

Results from the study-completer sample converged with those of the larger samples, as there was a main effect of development on SSRT [*F*_1,69_ = 4.383, *p* =.040, η^2^ = .065], where SSRT decreased over development. In addition, we found an age [*F*_1,36_ = 6.236, *p* = .017] and site effect [*F*_3,33_ = 4.975, *p* = .006] on MRT. We found no main effects or interaction effects of diagnostic group or development for any of the other behavioral measures (MRT, SSD, number of omissions, commission errors) in the study-completer sample (Table S4).

There were no correlations between developmental change in task performance and symptom severity as assessed with questionnaires (RBS-R, CY-BOCS, CPRS-R).

### 3.3 ROI analysis

Using LME, we found an interaction in the fMRI sample between diagnostic group and development for activity in right precentral gyrus (PreC) during failed-stop trials [*F*_2, 73_ = 5.644, *p* = .005], driven by increases in activity for the OCD group over development (t(89) = -2.865, *p* = .005), while activity in other groups remained stable over development (Fig 2). This finding was not replicated in the study-completer sample (N = 40).

**Fig 2.**
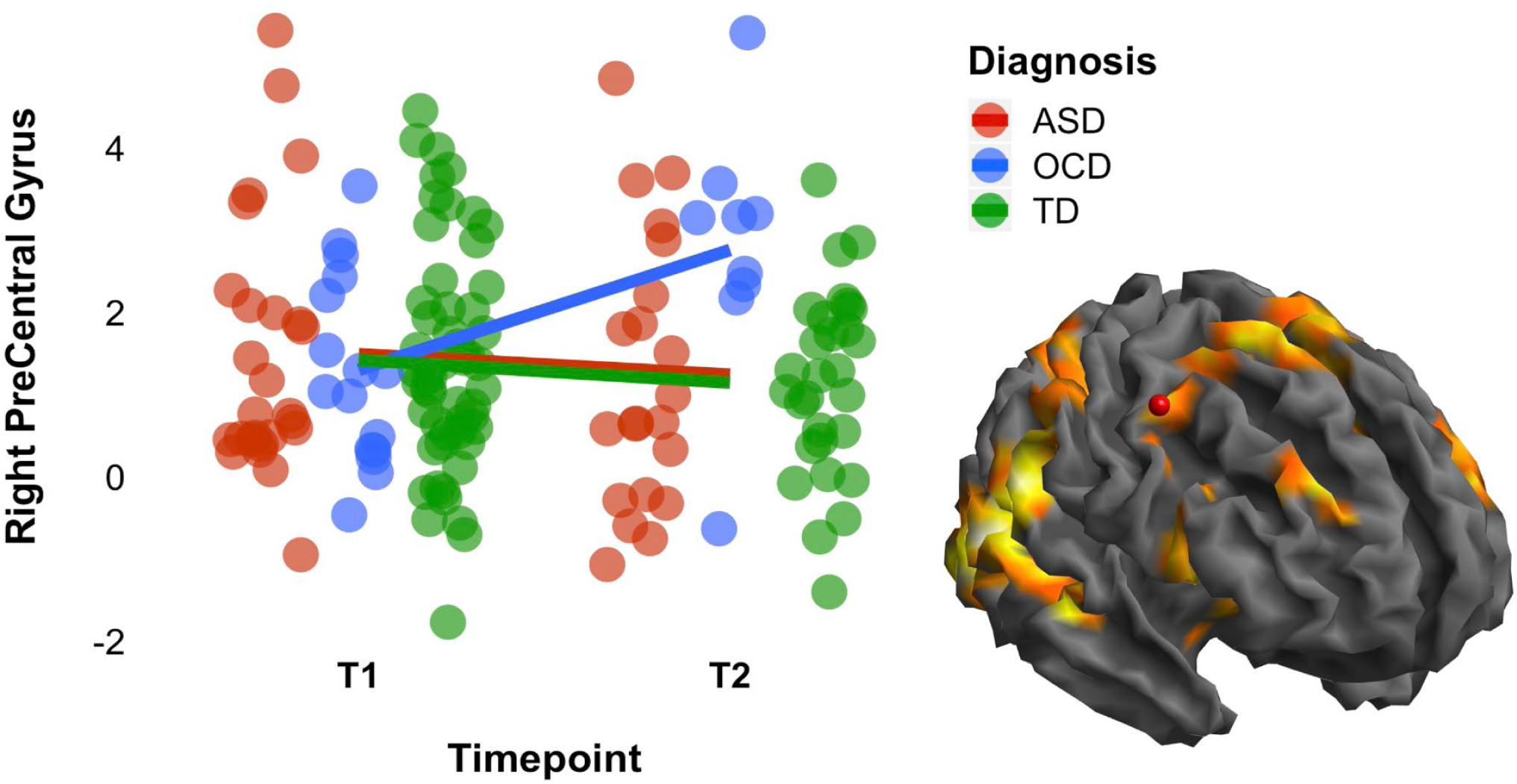
Activity in right precentral gyrus at baseline and follow-up during the fMRI stop-signal task. Lines represent average slopes. Activity increased with development for children with OCD during failed stopping. Note: the red dot in brain marks the right precentral gyrus area. Abbreviations: ASD, Autism Spectrum Disorder; OCD, Obsessive-Compulsive Disorder; TD, typically developing; T1, timepoint 1/baseline; T2, timepoint 2/follow-up

### 3.4 Whole-Brain effects

In line with our baseline results, we found no group differences in brain activation that survived whole-brain correction (*p*_FWE_ < .05). Nor were there any main effects of development, or group by development interactions in brain activity during successful and failed stopping.

### 3.5 Brain-behavior relationships

Brain-behavior correlation analyses showed that for typically developing children, change in activity in right middle cingulate gyrus, was positively associated with the developmental change in SSRT [*r* = 0.59, *p* = .007], and negatively with change in SSD [*r* = - 0.62, *p* = .005] during successful stop trials (Table S5A). During failed stop-trials, change in activity in right precentral gyrus correlated negatively with change in MRT [*r* = -0.79, *p* < .001] and change in SSD [*r* = -0.79, *p* < .001] (Table S5A).

We found no brain-behavior correlations in children with ASD and OCD (analyzed together) (Table S5B), nor did we find associations between change in task performance and change in brain activity in children with lower or higher ADHD symptoms that survived Bonferroni-correction.

## 4. Discussion

We examined the development of the behavioral and neural correlates of cognitive control in a multicenter, a little over a year longitudinal study, using a modified stop-signal task in children with ASD or OCD, and typically developing (TD) children. At baseline, children with ASD or OCD showed no changes in cognitive control or changes in brain activity in task-relevant neural networks compared to TD children (Gooskens et al., 2019). In the current study, we found the expected developmental improvement in cognitive control regardless of diagnostic group, with only subtle differences between the groups in terms of the development of behavior and brain activity. The results did not support our hypotheses on developmental changes in cognitive control and associated neural circuitry in ASD and OCD, compared to typical development. Nor did we find support for the hypothesis that cognitive control as assessed by the stop-signal task is associated with repetitive behavior in these children. Rather, our findings suggest that, similar to the lack of differences at baseline, the development of cognitive control in children with ASD or OCD follows a pattern similar to that of typically developing children, at least in the age range investigated here.

As expected, we found that SSRT decreased across groups with no statistically significant group differences in baseline or follow-up average scores, indicating improvement in cognitive control with development. The findings are in line with previous work in typical development (Huizinga et al., 2006). We found no evidence that children with ASD or OCD showed delayed development of cognitive control over this short time. As such, these findings support the suggestion from our earlier study that any impairments in cognitive control in ASD and OCD may emerge later during development, to support findings of decreased cognitive control in adults with OCD (Chamberlain et al., 2007; Kang et al., 2012; Marzuki et al., 2020; Penadés et al., 2007; de Wit et al., 2012). This raises questions about the place of cognitive control in a mechanistic, causal cascade of repetitive behavior in these disorders.

We also did not find the hypothesized delay in the development of prefrontal brain activity in children with ASD and OCD. Rather, we found a subtle developmental difference where activity in right precentral gyrus during failed-stop trials increased for children with OCD. The precentral gyrus is part of Brodmann area 6 (BA6), including pre-motor cortex and supplementary motor area (SMA), and is involved in the planning of movements. Activity in this region during response inhibition has previously been reported in healthy adults, as well as adults with OCD (Roth et al, 2007). Speculatively, this finding may suggest that, in contrast to children with ASD and TD children, children with OCD recruit precentral gyrus more for cognitive control during late childhood.

We found some evidence of associations between task performance and brain activity in typically developing children. A developmental increase in task performance (reflected by shorter SSRTs) was associated with decreased activity in right middle cingulate gyrus (MCG) during successful stopping. As opposed to typically developing children, we found no brain-behavior correlations in children with ASD and OCD (analyzed together). Also, we found no evidence that either task performance or activity in frontostriatal circuitry was related to severity of or changes in repetitive behavior in children with ASD and OCD, similar to what has been found cross-sectionally using similar tasks (Ambrosino et al. 2014). However, we cannot rule out that the relatively short interval (average 1.23 years) between both assessments and the reduction in symptoms with age may have affected this null-result.

It is not uncommon in ASD and OCD fMRI research to find differences in brain activation despite comparable task performance, and vice versa. Moreover, associations with clinical symptom ratings are frequently lacking (Wooley et al., 2008). Given the problems these children experience in everyday life, these observations raise the question to what extent cognitive control problems in ASD or OCD are fully captured by any single task we use in research (Geurts, Corbett and Solomon, 2009). It has been suggested that findings of deficits in cognitive control are dependent on the type of task, with diagnostic differences observed more frequently on interference tasks (Adam and Jarrold 2012; Christ, 2007) than stop-signal tasks (Adams and Jarrold, 2012; Chantiluke et al., 2015; Gooskens et al., 2019; Ozonoff and Strayer, 1997; Albajara Sáenz et al., 2020; Schmitt et al., 2017). Moreover, cognitive control may be affected only when particularly salient stimuli are used as stimuli

(Yerys et al., 2013; Bos et al., 2018; 2019). In line with this notion, cognitive control has been suggested to be a multi-faceted construct, related to distinct, but overlapping brain circuits, where distinct components of cortico-striatal-thalamocortical (CSTC) circuits may be related to different types of control and repetitive behavior (Carlisi et al., 2017; Friedmann and Miyake., 2004; Zhang et al., 2017; Nigg, 2017; Langen et al., 2011; Mataix-Cols et al., 2004). This may explain some of the inconsistency between tasks and studies.

The findings from our study should be considered in the context of some practical limitations and strengths. First, and inherent to longitudinal studies, there were drop-outs at follow-up which led to a modest final sample size, and may have affected our ability to find developmental effects. We addressed this issue by using two different analysis strategies: an intent-to-treat analysis by means of linear mixed-effects modelling to reduce bias introduced by drop-outs, and a study-completer analysis by means of repeated measures ANOVA design. Furthermore, although this longitudinal study with two measurements already provides valuable information on the development of cognitive control, studies with three or more timepoints would permit the more precise mapping of trajectories.

To summarize, we found only subtle differences in the development of cognitive control and associated brain circuitry in children with OCD. We found no notable differences in cognitive control or brain activity in children with ASD or OCD compared to TD children. We found no evidence that cognitive control, as assessed by the stop-signal task, was associated with repetitive behavior in children with ASD and OCD. Heterogeneity of samples (including in age) and dissimilarity in task design are factors that likely contribute to the inconsistency in findings between studies. Therefore, to detect differences in these neurodevelopmental disorders it will be critical to run longitudinal studies with larger sample sizes and more than two timepoints and with longer time-intervals, using similar task designs, and to adopt a dimensional approach to mapping cognitive control.

## Acknowledgements

We would like to thank the participants and their parents for participating in this study. Further, we would like to thank Vincent Mensen, Devon Shook, Saskia de Ruiter and Isabella Wolf for their assistance in data collection and analysis. Steven Williams would like to thank the National Institute for Health Research (NIHR) Biomedical Research Centre at South London and Maudsley NHS Foundation Trust and King’s College London for their support.

## Ethical approval

The study was approved by local ethics committees for each site (Nijmegen and Utrecht: Commissie Mensgebonden Onderzoek Regio Arnhem-Nijmegen, 2013, NL nr: 42004.091.12; Mannheim: Ethics committee of the Medical Faculty Mannheim, Heidelberg University, 2013, nr: 213-616 N-MA; London: NRES Committee London - Camberwell St Giles, 2013, nr: 14/LO/1413).

## References

Adams, N.C., Jarrold, C., 2012. Inhibition in autism: Children with autism have difficulty inhibiting irrelevant distractors but not prepotent responses. J. Autism Dev. Disord. 42, 1052–1063. https://doi.org/10.1007/s10803-011-1345-3

Albajara Sáenz, A., Septier, M., Schuerbeek, P. Van, Baijot, S., Deconinck, N., Defresne, P., Delvenne, V., Passeri, G., Raeymaekers, H., Salvesen, L., Victoor, L., Villemonteix, T., Willaye, E., Peigneux, P., Massat, I., n.d. ADHD and ASD: distinct brain patterns of inhibition-related activation? https://doi.org/10.1038/s41398-020-0707-z

Ambrosino, S., Bos, D.J., Van Raalten, T.R., Kobussen, N.A., Van Belle, J., Oranje, B., Durston, S., 2014. Functional connectivity during cognitive control in children with autism spectrum disorder: An independent component analysis. J. Neural Transm. 121, 1145–1155. https://doi.org/10.1007/s00702-014-1237-8

American Psychiatric Association. (2000). Diagnostic and statistical manual of mental disorders (4th ed., text rev.). Washington, DC: Author.

American Psychiatric Association. (2013). Diagnostic and statistical manual of mental disorders (5th ed.). Washington, DC: Author

Bates D., Maechler, M., Bolker, B., Walker, S. (2015). Fitting Linear Mixed-Effects Models Using lme4. Journal of Statistical Software, 67(1), 1–48. doi:10.18637/jss.v067.i01.

Bodfish, J.W., Symons, F.J., Parker, D.E., Lewis, M.H., 2000. Varieties of repetitive behavior in autism: Comparisons to mental retardation. J. Autism Dev. Disord. 30, 237–243. https://doi.org/10.1023/A:1005596502855

Bos, D.J., Silver, B.M., Barnes, E.D., Ajodan, E.L., Silverman, M.R., Clark-Whitney, E., Tarpey, T., Jones, R.M., 1234. Adolescent-Specific Motivation Deficits in Autism Versus Typical Development. J. Autism Dev. Disord. 50, 364–372. https://doi.org/10.1007/s10803-019-04258-9

Bos, D.J., Silverman, M.R., Ajodan, E.L., Martin, C., Silver, B.M., Brouwer, G.J., Di Martino, A., Jones, R.M., 2019. Rigidity coincides with reduced cognitive control to affective cues in children with autism. J. Abnorm. Psychol. 128, 431–441. https://doi.org/10.1037/abn0000423

Bunge, S.A., Dudukovic, N.M., Thomason, M.E., Vaidya, C.J., Gabrieli, J.D.E., 2002. Immature frontal lobe contributions to cognitive control in children: Evidence from fMRI. Neuron 33, 301–311. https://doi.org/10.1016/S0896-6273(01)00583-9

Carlisi, C.O., Norman, L.J., Lukito, S.S., Radua, J., Mataix-Cols, D., Rubia, K., 2017. Comparative Multimodal Meta-analysis of Structural and Functional Brain Abnormalities in Autism Spectrum Disorder and Obsessive-Compulsive Disorder. Biol. Psychiatry 82, 83–102. https://doi.org/10.1016/j.biopsych.2016.10.006

Chamberlain, S.R., Blackwell, A.D., Fineberg, N.A., Robbins, T.W., Sahakian, B.J., 2005. The neuropsychology of obsessive compulsive disorder: The importance of failures in cognitive and behavioural inhibition as candidate endophenotypic markers. Neurosci. Biobehav. Rev. 29, 399–419. https://doi.org/10.1016/j.neubiorev.2004.11.006

Chamberlain, S.R., Fineberg, N.A., Menzies, L.A., Blackwell, A.D., Bullmore, E.T., Robbins, T.W., Sahakian, B.J., 2007. Impaired cognitive flexibility and motor inhibition in unaffected first-degree relatives of patients with obsessive-compulsive disorder. Am. J. Psychiatry 164, 335–338. https://doi.org/10.1176/ajp.2007.164.2.335

Christ, S.E., Holt, D.D., White, D.A., Green, L., 2007. Inhibitory control in children with autism spectrum disorder. J. Autism Dev. Disord. 37, 1155–1165. https://doi.org/10.1007/s10803-006-0259-y

Cohen, J.R., Asarnow, R.F., Sabb, F.W., Bilder, R.M., Bookheimer, S.Y., Knowlton, B.J., Poldrack, R.A., 2010. Decoding developmental differences and individual variability in response inhibition through predictive analyses across individuals. Front. Hum. Neurosci. 4, 1–12. https://doi.org/10.3389/fnhum.2010.00047

De Wit, S.J., De Vries, F.E., Van Der Werf, Y.D., Cath, D.C., Heslenfeld, D.J., Veltman, E.M., Van Balkom, A.J.L.M., Veltman, D.J., Van Den Heuvel, O.A., 2012. Presupplementary motor area hyperactivity during response inhibition: A candidate endophenotype of obsessive-compulsive disorder. Am. J. Psychiatry 169, 1100–1108. https://doi.org/10.1176/appi.ajp.2012.12010073

Durston, S., Davidson, M.C., Tottenham, N., Galvan, A., Spicer, J., Fossella, J.A., Casey, B.J., 2006. A shift from diffuse to focal cortical activity with development: Commentary. Dev. Sci. 9, 1–20. https://doi.org/10.1111/j.1467-7687.2005.00454.x

Durston, S., Nederveen, H., Van Dijk, S., Van Belle, J., De Zeeuw, P., Langen, M., Van Dijk, A., 2009. Magnetic resonance simulation is effective in reducing anxiety related to magnetic resonance scanning in children. J. Am. Acad. Child Adolesc. Psychiatry. https://doi.org/10.1097/CHI.0b013e3181930673

Durston, S., Thomas, K.M., Yang, Y., Uluǧ, A.M., Zimmerman, R.D., Casey, B.J., 2002. A neural basis for the development of inhibitory control. Dev. Sci. 5, 9–16. https://doi.org/10.1111/1467-7687.00235

Friedman, N.P., Miyake, A., 2004. The Relations Among Inhibition and Interference Control Functions: A Latent-Variable Analysis. J. Exp. Psychol. Gen. 133, 101–135. https://doi.org/10.1037/0096-3445.133.1.101

Geurts, H.M., Corbett, B., Solomon, M., 2009. The paradox of cognitive flexibility in autism. Trends Cogn. Sci. https://doi.org/10.1016/j.tics.2008.11.006

Goodman, R., Ford, T., Richards, H., Gatward, R., Meltzer, H., 2000. The Development and Well-Being Assessment: Description and Initial Validation of an Integrated Assessment of Child and Adolescent Psychopathology. J. Child Psychol. Psychiatry 41, 645–655. https://doi.org/10.1111/j.1469-7610.2000.tb02345.x

Gooskens, B., Bos, D.J., Mensen, V.T., Shook, D.A., Bruchhage, M.M.K., Naaijen, J., Wolf, I., Brandeis, D., Williams, S.C.R., Buitelaar, J.K., Oranje, B., Durston, S., 2019. No evidence of differences in cognitive control in children with autism spectrum disorder or obsessive-compulsive disorder: An fMRI study. Dev. Cogn. Neurosci. 36. https://doi.org/10.1016/j.dcn.2018.11.004

Hill, E.L., 2004. Executive dysfunction in autism. Trends Cogn. Sci. 8, 26–32. https://doi.org/10.1016/j.tics.2003.11.003

Huizinga, M., Dolan, C. V., van der Molen, M.W., 2006. Age-related change in executive function: Developmental trends and a latent variable analysis. Neuropsychologia 44, 2017–2036. https://doi.org/10.1016/j.neuropsychologia.2006.01.010

IBM SPSS Statistics for Windows, version XX (IBM Corp., Armonk, N.Y., USA)

Ivarsson, T., Melin, K., 2008. Autism spectrum traits in children and adolescents with obsessive-compulsive disorder (OCD). J. Anxiety Disord. 22, 969–978. https://doi.org/10.1016/j.janxdis.2007.10.003

Jiujias, M., Kelley, E., Hall, L., 2017. Restricted, Repetitive Behaviors in Autism Spectrum Disorder and Obsessive–Compulsive Disorder: A Comparative Review. Child Psychiatry Hum. Dev. 48, 944–959. https://doi.org/10.1007/s10578-017-0717-0

Kang, D.H., Jang, J.H., Han, J.Y., Kim, J.H., Jung, W.H., Choi, J.S., Choi, C.H., Kwon, J.S., 2013. Neural correlates of altered response inhibition and dysfunctional connectivity at rest in obsessive-compulsive disorder. Prog. Neuro-Psychopharmacology Biol. Psychiatry 40, 340–346. https://doi.org/10.1016/j.pnpbp.2012.11.001

Kaufman, J., Birmaher, B., Brent, D., Rao, U., Flynn, C., Moreci, P., Williamson, D., Ryan, N., 1997. Schedule for affective disorders and schizophrenia for school-age children-present and lifetime version (K-SADS-PL): Initial reliability and validity data. J. Am. Acad. Child Adolesc. Psychiatry 36, 980–988. https://doi.org/10.1097/00004583-199707000-00021

Keith Conners, C., Sitarenios, G., Parker, J.D.A., Epstein, J.N., 1998. The revised Conners’ Parent Rating Scale (CPRS-R): Factor structure, reliability, and criterion validity. J. Abnorm. Child Psychol. 26, 257–268. https://doi.org/10.1023/A:1022602400621

Langen, M., Durston, S., Kas, M.J.H., van Engeland, H., Staal, W.G., 2011. The neurobiology of repetitive behavior: …and men. Neurosci. Biobehav. Rev. https://doi.org/10.1016/j.neubiorev.2010.02.005

Leyfer, O.T., Folstein, S.E., Bacalman, S., Davis, N.O., Dinh, E., Morgan, J., Tager-Flusberg, H., Lainhart, J.E., 2006. Comorbid psychiatric disorders in children with autism: Interview development and rates of disorders. J. Autism Dev. Disord. 36, 849–861. https://doi.org/10.1007/s10803-006-0123-0

Lord, C., Rutter, M., Le Couteur, A., 1994. Autism Diagnostic Interview-Revised: A revised version of a diagnostic interview for caregivers of individuals with possible pervasive developmental disorders. J. Autism Dev. Disord. 24, 659–685. https://doi.org/10.1007/BF02172145

Luna, B., Doll, S.K., Hegedus, S.J., Minshew, N.J., Sweeney, J.A., 2007. Maturation of Executive Function in Autism. Biol. Psychiatry 61, 474–481. https://doi.org/10.1016/j.biopsych.2006.02.030

Luna, B., Marek, S., Larsen, B., Tervo-Clemmens, B., Chahal, R., 2015. An Integrative Model of the Maturation of Cognitive Control. Annu. Rev. Neurosci. 38, 151–170. https://doi.org/10.1146/annurev-neuro-071714-034054

Luna, B., Padmanabhan, A., O’Hearn, K., 2010. What has fMRI told us about the Development of Cognitive Control through Adolescence? Brain Cogn. https://doi.org/10.1016/j.bandc.2009.08.005

Marzuki, A.A., Pereira de Souza, A.M.F.L., Sahakian, B.J., Robbins, T.W., 2020. Are candidate neurocognitive endophenotypes of OCD present in paediatric patients? A systematic review. Neurosci. Biobehav. Rev. 108, 617–645. https://doi.org/10.1016/j.neubiorev.2019.12.010

Mataix-Cols, D., Wooderson, S., Lawrence, N., Brammer, M.J., Speckens, A., Phillips, M.L., 2004. Distinct neural correlates of washing, checking, and hoarding symptom dimensions in obsessive-compulsive disorder. Arch. Gen. Psychiatry 61, 564–576. https://doi.org/10.1001/archpsyc.61.6.564

McDougle, C.J., Kresch, L.E., Goodman, W.K., Naylor, S.T., Volkmar, F.R., Cohen, D.J., Price, L.H., 1995. A case-controlled study of repetitive thoughts and behavior in adults with autistic disorder and obsessive-compulsive disorder. Am. J. Psychiatry 152, 772–777. https://doi.org/10.1176/ajp.152.5.772

Moritz, S., Birkner, C., Kloss, M., Jahn, H., Hand, I., Haasen, C., Krausz, M., 2002. Executive functioning in obsessive-compulsive disorder, unipolar depression, and schizophrenia. Arch. Clin. Neuropsychol. 17, 477–483. https://doi.org/10.1016/S0887-6177(01)00130-5

Mosconi, M.W., Kay, M., D’Cruz, A.M., Seidenfeld, A., Guter, S., Stanford, L.D., Sweeney, J.A., 2009. Impaired inhibitory control is associated with higher-order repetitive behaviors in autism spectrum disorders. Psychol. Med. 39, 1559–1566. https://doi.org/10.1017/S0033291708004984

Naaijen, J., de Ruiter, S., Zwiers, M.P., Glennon, J.C., Durston, S., Lythgoe, D.J., Williams, S.C.R., Banaschewski, T., Brandeis, D., Franke, B., Buitelaar, J.K., 2016. COMPULS: Design of a multicenter phenotypic, cognitive, genetic, and magnetic resonance imaging study in children with compulsive syndromes. BMC Psychiatry 16. https://doi.org/10.1186/s12888-016-1072-6

Nigg, J.T., 2017. Annual Research Review: On the relations among self-regulation, selfcontrol, executive functioning, effortful control, cognitive control, impulsivity, risk-taking, and inhibition for developmental psychopathology. J. Child Psychol. Psychiatry Allied Discip. 58, 361–383. https://doi.org/10.1111/jcpp.12675

Penadés, R., Catalán, R., Rubia, K., Andrés, S., Salamero, M., Gastó, C., 2007. Impaired response inhibition in obsessive compulsive disorder. Eur. Psychiatry 22, 404–410. https://doi.org/10.1016/j.eurpsy.2006.05.001

R Core Team (2019). R: A language and environment for statistical computing. R Foundation for Statistical Computing, Vienna, Austria. URL https://www.R-project.org/

Rubia, K., Cubillo, A., Smith, A.B., Woolley, J., Heyman, I., Brammer, M.J., 2010. Disorder-specific dysfunction in right inferior prefrontal cortex during two inhibition tasks in boys with attention-deficit hyperactivity disorder compared to boys with obsessivecompulsive disorder. Hum. Brain Mapp. 31, 287–299. https://doi.org/10.1002/hbm.20864

Rubia, K., Smith, A.B., Brammer, M.J., Taylor, E., 2003. Right inferior prefrontal cortex mediates response inhibition while mesial prefrontal cortex is responsible for error detection. Neuroimage 20, 351–358. https://doi.org/10.1016/S1053-8119(03)00275-1

Scahill, L., Riddle, M.A., McSwiggin-Hardin, M., Ort, S.I., King, R.A., Goodman, W.K., Cicchetti, D., Leckman, J.F., 1997. Children’s Yale-Brown Obsessive Compulsive Scale: Reliability and validity. J. Am. Acad. Child Adolesc. Psychiatry 36, 844–852. https://doi.org/10.1097/00004583-199706000-00023

Schmitt, L.M., White, S.P., Cook, E.H., Sweeney, J.A., Mosconi, M.W., 2018. Cognitive mechanisms of inhibitory control deficits in autism spectrum disorder. J. Child Psychol. Psychiatry Allied Discip. 59, 586–595. https://doi.org/10.1111/jcpp.12837

Shaffer, D., Fisher, P., Lucas, C.P., Dulcan, M.K., Schwab-Stone, M.E., 2000. NIMH Diagnostic Interview Schedule for Children Version IV (NIMH DISC-IV): Description, differences from previous versions, and reliability of some common diagnoses. J. Am. Acad. Child Adolesc. Psychiatry 39, 28–38. https://doi.org/10.1097/00004583-200001000-00014

Snyder, H.R., Kaiser, R.H., Warren, S.L., Heller, W., 2015. Obsessive-Compulsive Disorder Is Associated With Broad Impairments in Executive Function: A Meta-Analysis. Clin. Psychol. Sci. https://doi.org/10.1177/2167702614534210

Solomon, M., Ozonoff, S.J., Cummings, N., Carter, C.S., 2008. Cognitive control in autism spectrum disorders. Int. J. Dev. Neurosci. 26, 239–247. https://doi.org/10.1016/j.ijdevneu.2007.11.001

Somerville, L.H., Hare, T., Casey, B., 2008. This model, consistent with others. J Cogn Neurosci 23, 2123–2134. https://doi.org/10.1162/jocn.2010.21572

Tamm, L., Menon, V., Reiss, A.L., 2002. Maturation of Brain Function Associated with Response Inhibition. J. Am. Acad. Child Adolesc. Psychiatry 41, 1231–1238. https://doi.org/10.1097/00004583-200210000-00013

Verbruggen, F., Aron, A.R., Band, G.P.H., Beste, C., Bissett, P.G., Brockett, A.T., Brown, J.W., Chamberlain, S.R., Chambers, C.D., Colonius, H., Colzato, L.S., Corneil, B.D., Coxon, J.P., Dupuis, A., Eagle, D.M., Garavan, H., Greenhouse, I., Heathcote, A., Huster, R.J., Jahfari, S., Kenemans, J.L., Leunissen, I., Li, C.S.R., Logan, G.D., Matzke, D., Morein-Zamir, S., Murthy, A., Paré, M., Poldrack, R.A., Ridderinkhof, K.R., Robbins, T.W., Roesch, M., Rubia, K., Schachar, R.J., Schall, J.D., Stock, A.K., Swann, N.C., Thakkar, K.N., Van Der Molen, M.W., Vermeylen, L., Vink, M., Wessel, J.R., Whelan, R., Zandbelt, B.B., Boehler, C.N., 2019. A consensus guide to capturing the ability to inhibit actions and impulsive behaviors in the stop-signal task. Elife 8. https://doi.org/10.7554/eLife.46323

Verbruggen, F., Chambers, C.D., Logan, G.D., 2013. Fictitious inhibitory differences: how skewness and slowing distort the estimation of stopping latencies. Psychol. Sci. 24, 352–62. https://doi.org/10.1177/0956797612457390

Woolley, J., Heyman, I., Brammer, M., Frampton, I., McGuire, P.K., Rubia, K., 2008. Brain activation in paediatric obsessive-compulsive disorder during tasks of inhibitory control. Br. J. Psychiatry 192, 25–31. https://doi.org/10.1192/bjp.bp.107.036558

Yerys, B.E., 2015. An Update on the Neurobiology of Repetitive Behaviors in Autism, International Review of Research in Developmental Disabilities. https://doi.org/10.1016/bs.irrdd.2015.06.006

Zandt, F., Prior, M., Kyrios, M., 2007. Repetitive behaviour in children with high functioning autism and obsessive compulsive disorder. J. Autism Dev. Disord. 37, 251–259. https://doi.org/10.1007/s10803-006-0158-2

Zandt, F., Prior, M., Kyrios, M., 2009. Similarities and differences between children and adolescents with autism spectrum disorder and those with obsessive compulsive disorder: Executive functioning and repetitive behaviour. Autism 13, 43–57. https://doi.org/10.1177/1362361308097120

Zhang, R., Geng, X., Lee, T.M.C., 2017. Large-scale functional neural network correlates of response inhibition: an fMRI meta-analysis. Brain Struct. Funct. 222, 3973–3990. https://doi.org/10.1007/s00429-017-1443-x

