## Supplemental Material for "The Development of Cognitive Control in Children with Autism Spectrum Disorders or Obsessive-Compulsive Disorder: A Longitudinal fMRI study"

**TABLE S1. SAMPLE CHARACTERISTICS [FULL FMRI SAMPLE]**

| | ASD | OCD | TD | Statistical test | *P*-value |
|---|---|---|---|---|---|
| ***N* T1** | 26 | 16 | 53 | | |
| ***N* T2** | 21 | 9 | 29 | | |
| **SEX (M/F) T1** | 17/9 | 7/9 | 29/24 | $\chi^2(2) = 1.936$ | .380 |
| **SEX (M/F) T2** | 14/7 | 5/4 | 16/13 | $\chi^2(2) = 0.729$ | .694 |
| **AGE IN YEARS T1, MEAN (SD)** | 11.33 (1.07) | 10.92 (1.47) | 10.76 (1.15) | $F_{2, 92} = 2.012$ | .140 |
| **AGE IN YEARS T2, MEAN (SD)** | 12.33 (1.56) | 12.43 (1.58) | 11.95 (1.27) | $F_{2, 92} = 1.290$ | .533 |
| **INTERVAL IN YEARS** | 1.00 | 1.51 | 1.19 | | |
| **ESTIMATED IQ T1, MEAN (SD)** | 108.88 (16.67) | 100.72 (13.26) | 111.93 (10.36) | K-W $\chi^2$ (2) = 7.999 | .018* |
| **ESTIMATED IQ T2, MEAN (SD)** | 110.87 (14.16) | 105.82 (11.21) | 115.03 (7.72) | K-W $\chi^2$ (2) = 3.515 | .172 |

Abbreviations: T1, Timepoint 1, T2, Timepoint 2, SD, standard deviation

**Table S2. Clinical and performance measures [full fMRI Sample]**

| | T1 ASD $N$ = 26 | T2 ASD $N$ = 21 | T1 OCD $N$ = 16 | T2 OCD $N$ = 9 | T1 TD $N$ = 53 | T2 TD $N$ = 29 | Statistics: Time | Statistics: Group | Statistics: Time* Group |
|---|---|---|---|---|---|---|---|---|---|
| | Mean (SD) | Mean (SD) | Mean (SD) | Mean (SD) | Mean (SD) | Mean (SD) | *P*-value | *P*-value | *P*-value |
| **Questionnaires** | | | | | | | | | |
| *CY-BOCS* | | | | | | | | | |
| - Obsessions | | | 7.19 (5.23) | 8.13 (6.24) | | | .124 | n.a. | n.a. |
| - Compulsions | | | 10.19 (3.51) | 7.78 (5.65) | | | .066 | n.a. | n.a. |
| - Total score | | | 17.38 (7.70) | 15.00 (11.74) | | | .038*[a] | n.a. | n.a. |
| *RBS-Revised* | | | | | | | | | |
| - Compulsivity | 2.04 (2.65) | 2.24 (2.84) | 4.69 (3.03) | 2.67 (2.12) | 0.06 (0.31) | 0.04 (0.19) | 0.273 | < .001*[f] | .041*[a] |
| - Total score | 20.04 (16.04) | 17.57 (17.37) | 16.47 (11.36) | 12.00 (10.25) | 0.64 (1.30) | 0.54 (1.14) | 0.319 | < .001*[f] | .030*[b] |
| *CPRS-Revised: Long* | | | | | | | | | |
| - Inattention | 62.85 (11.93) | 59.86 (10.16) | 57.36 (10.62) | 54.13 (8.31) | 45.63 (6.25) | 46.18 (6.19) | .388 | < .001*[f] | n.s. |
| - Hyperactivity | 62.90 (13.16) | 61.50 (12.67) | 61.29 (10.51) | 58.00 (10.62) | 46.92 (3.78) | 45.81 (4.24) | .959 | < .001*[f] | n.s. |
| - Total score | 64.40 (12.78) | 61.65 (11.03) | 59.93 (10.77) | 56.50 (6.89) | 45.26 (4.76) | 44.82 (4.11) | .256 | < .001*[f] | n.s. |
| **Performance** | | | | | | | | | |
| MRT | 561.27 (101.11) | 503.93 (67.28) | 516.08 (93.15) | 571.60 (113.23) | 516.41 (90.59) | 546.20 (56.39) | .112 | .669 | .006*[c] |
| SSD | 370.36 (87.36) | 331.99 (79.56) | 336.01(111.96) | 417.63 (114.26) | 381.72 (144.27) | 348.30 (56.56) | .500 | .695 | .049*[d] |
| SSRT | 165.94 (71.68) | 145.25 (51.48) | 190.72 (79.99) | 120.65 (61.30) | 188.67 (69.32) | 168.89 (63.87) | .047*[e] | .423 | n.s. |
| Omissions | 2.02 % | 2.20 % | 2.43 % | 1.66 % | 2.24 % | 1.58 % | .934 | .865 | n.s. |
| Comissions | 2.79 % | 3.63 % | 4.25 % | 3.56 % | 4.37 % | 3.99 % | .351 | .529 | n.s. |
| Successful Stopping | 52.4 % | 51.4 % | 51.2 % | 52.8 % | 51.41% | 51.38 % | .695 | .522 | n.s. |

Abbreviations: T1, Timepoint 1, T2, Timepoint 2, ASD, autism spectrum disorder; OCD, obsessive-compulsive disorder; TD, typically developing group; SD, standard deviation; m/f, male/female; ADI, Autism Diagnostic Interview; CY-BOCS, Children's Yale-Brown Obsessive-Compulsive scale; RBS, Repetitive-Behavior scale; CPRS, Conners' Parent Rating scale, MRT = Mean reaction time, SSD = Stop-signal delay, SSRT = Stop-signal reaction time, n.a. not applicable, n.s. not significant (removed from model)

[a] Children with OCD show a decrease in compulsive behavior over development

[b] Children with ASD show a decrease in total severity of repetitive behavior over development

[c] Children with OCD and TD show slower reaction times over development

[d] Children with OCD show greater SSD-latencies over development

[e] All children show decreased SSRTs over development

[f] ASD, OCD > TD

**TABLE S3. SAMPLE CHARACTERISTICS [STUDY COMPLETER SAMPLE]**

| | ASD | OCD | TD | Statistical test | *P*-value |
|---|---|---|---|---|---|
| ***N*** | 14 | 6 | 20 | | |
| **SEX (M/F)** | 8/6 | 3/3 | 7/13 | $\chi^2(2) = 1.703$ | .202 |
| **AGE IN YEARS T1, MEAN (SD)** | 11.39 (1.37) | 10.34 (1.25) | 10.43 (1.18) | $F_{2, 37} = 2.783$ | .075 |
| **AGE IN YEARS T2, MEAN (SD)** | 12.89 (1.36) | 11.73 (1.47) | 11.92 (1.27) | $F_{2, 37} = 2.709$ | .080 |
| **INTERVAL IN YEARS** | 1.5 | 1.39 | 1.49 | | |
| **ESTIMATED IQ T1, MEAN (SD)** | 110.46 (16.99) | 100.41 (6.94) | 115.32 (7.80) | K-W $\chi^2$ (2) = 8.740 | .013* |
| **ESTIMATED IQ T2, MEAN (SD)** | 108.59 (14.37) | 99.49 (7.03) | 114.87 (7.94) | $F_{2, 37} = 5.191$ | .010* |

Abbreviations: T1, Timepoint 1, T2, Timepoint 2, SD, standard deviation

**Table S4. Clinical and performance measures [study completer sample]**

| | T1 ASD *N* = 14 | T2 ASD *N* = 14 | T1 OCD *N* = 6 | T2 OCD *N* = 6 | T1 TD *N* = 20 | T2 TD *N* = 20 | Statistics | | |
|---|---|---|---|---|---|---|---|---|---|
| | | | | | | | Time | Group | Time* Group |
| | Mean (SD) | Mean (SD) | Mean (SD) | Mean (SD) | Mean (SD) | Mean (SD) | *P*-value | *P*-value | *P*-value |
| **Questionnaires** | | | | | | | | | |
| *CY-BOCS* | | | | | | | | | |
| - Obsessions | | | 6.83 (5.78) | 6.60 (6.27) | | | .080 | n.a. | n.a. |
| - Compulsions | | | 10.17 (3.19) | 6.67 (5.43) | | | .022*[a] | n.a. | n.a. |
| - Total score | | | 17.00 (8.39) | 12.17 (11.13) | | | .025*[a] | n.a. | n.a. |
| *RBS-Revised* | | | | | | | | | |
| - Compulsivity | 1.50 (2.31) | 1.64 (1.82) | 3.83 (3.49) | 2.33 (2.07) | 0.15 (0.49) | 0.05 (0.22) | 0.884 | < .001*[d] | n.s. |
| - Total score | 18.57 (14.81) | 13.64 (11.51) | 13.83 (13.05) | 13.00 (11.31) | 0.60 (1.54) | 0.65 (1.27) | 0.251 | < .001*[d] | .014*[b] |
| *CPRS-Revised: Long* | | | | | | | | | |
| - Inattention | 62.09 (13.09) | 61.36 (11.13) | 55.80 (12.32) | 50.80 (4.09) | 47.17 (8.10) | 47.40 (6.86) | .388 | < .001*[e] | n.s. |
| - Hyperactivity | 61.60 (13.68) | 60.15 (14.02) | 61.60 (11.68) | 57.60 (9.69) | 47.09 (4.76) | 46.79 (4.71) | .959 | < .001*[d] | n.s. |
| - Total score | 63.20 (13.48) | 62.08 (12.86) | 59.00 (12.35) | 54.20 (5.54) | 45.91 (4.97) | 45.68 (4.46) | .256 | < .001*[d] | n.s. |
| **Performance** | | | | | | | | | |
| MRT | 567.35 (67.59) | 499.18 (58.57) | 576.49 (53.45) | 551.39 (79.15) | 584.48 (91.69) | 549.42 (55.38) | .278 | .519 | n.s. |
| SSD | 384.17 (92.36) | 328.78 (85.89) | 361.73 (63.39) | 403.14 (123.76) | 345.26 (102.39) | 348.74 (58.07) | .553 | .461 | n.s. |
| SSRT | 146.78 (65.72) | 143.89 (55.34) | 178.50 (58.32) | 115.92 (70.87) | 201.85 (60.09) | 170.77 (61.36) | .040*[c] | .053 | n.s. |
| Omissions | 2.14 % | 2.38 % | 2.92 % | 2.14 % | 2.46 % | 2.07 % | .459 | .853 | n.s. |
| Comissions | 1.74 % | 2.44 % | 4.70 % | 4.77 % | 4.51 % | 4.12 % | .381 | .342 | n.s. |
| Successful Stopping | 52.6 % | 51.7 % | 52.78 % | 51.95 % | 51.67 % | 51.42 % | .991 | .406 | n.s. |

Abbreviations: T1, Timepoint 1, T2, Timepoint 2, ASD, autism spectrum disorder; OCD, obsessive-compulsive disorder; TD, typically developing group; SD, standard deviation; m/f, male/female; ADI, Autism Diagnostic Interview; CY-BOCS, Children's Yale-Brown Obsessive-Compulsive scale; RBS, Repetitive-Behavior scale; CPRS, Conners' Parent Rating scale, MRT = Mean reaction time, SSD = Stop-signal delay, SSRT = Stop-signal reaction time, n.a. not applicable, n.s. not significant (removed from model)

[a] Children with OCD show a decrease in compulsive behavior over time

[b] Children with ASD show a decrease in total severity of repetitive behavior over time

[c] All children show decreased SSRTs over development

[d] ASD, OCD > TD

[e] ASD > TD

**Table S5A. Spearman's correlations between ROIs and task performance measures in typical developing children of the study completer sample**

| Successful Stopping | Left Hemisphere | | | | Right Hemisphere | | |
|---|---|---|---|---|---|---|---|
| | Insula | MCG | MFG | SFG | Insula | MCG | MFG |
| **MRT** | .16 | .27 | .028* | .07 | .61 | .013* | .35 |
| **SSD** | .21 | .50 | .19 | .06 | .69 | **.005**** | .27 |
| **SSRT** | .76 | .24 | .72 | .52 | .92 | **.007**** | .030* |

| Failed Stopping | Left Hemisphere | | | Right Hemisphere | | | | |
|---|---|---|---|---|---|---|---|---|
| | Insula | MCG | MFG | Insula | PreC | SMA | MCG | MFG |
| **MRT** | .20 | .19 | .11 | .49 | **< .001**** | .030* | .047* | .14 |
| **SSD** | .50 | .19 | .016* | .48 | **< .001**** | .008* | .024* | .13 |
| **SSRT** | .68 | .019* | .014* | .89 | .012 | .033* | .14 | .30 |

Note: numbers are *p*-values; * = significant uncorrected ($p < .05$); ** = significant after Bonferroni correction (in bold font). Abbreviations: MCG = middle cingulate gyrus, MFG = middle frontal gyrus, SFG = superior frontal gyrus, PreC = precentral gyrus, SMA = supplementary motor area.

**Table S5B. Spearman's correlations between ROIs and task performance measures in patients (children with ASD and OCD analyzed together) of the study completer sample**

| Successful Stopping | Left Hemisphere | | | | Right Hemisphere | | |
|---|---|---|---|---|---|---|---|
| | Insula | MCG | MFG | SFG | Insula | MCG | MFG |
| **MRT** | .50 | .41 | .16 | .11 | .026* | .57 | .22 |
| **SSD** | .79 | .15 | .09 | .07 | .30 | .14 | .30 |
| **SSRT** | .82 | .21 | .040* | .18 | .91 | .18 | .63 |

| Failed Stopping | Left Hemisphere | | | Right Hemisphere | | | | |
|---|---|---|---|---|---|---|---|---|
| | Insula | MCG | MFG | Insula | PreC | SMA | MCG | MFG |
| **MRT** | .57 | .75 | .30 | .06 | .37 | .87 | .97 | .16 |
| **SSD** | .47 | .17 | .82 | .030* | .09 | .57 | .26 | .29 |
| **SSRT** | .60 | .19 | .95 | .13 | .24 | .91 | .12 | .79 |

Note: numbers are *p*-values, * = significant uncorrected ($p < .05$)
Abbreviations: MCG = middle cingulate gyrus, MFG = middle frontal gyrus, SFG = superior frontal gyrus, PreC = precentral gyrus, SMA = supplementary motor area
